# FuncFetch: An LLM-assisted workflow enables mining thousands of enzyme-substrate interactions from published manuscripts

**DOI:** 10.1101/2024.07.22.604620

**Authors:** Nathaniel Smith, Xinyu Yuan, Chesney Melissinos, Gaurav Moghe

**Affiliations:** Plant Biology Section, School of Integrative Plant Science, Cornell University, Ithaca, NY, USA 14853

## Abstract

**Motivation:** Thousands of genomes are publicly available, however, most genes in those genomes have poorly defined functions. This is partly due to a gap between previously published, experimentally-characterized protein activities and activities deposited in databases. This activity deposition is bottlenecked by the time-consuming biocuration process. The emergence of large language models (LLMs) presents an opportunity to speed up text-mining of protein activities for biocuration.

**Results:** We developed FuncFetch — a workflow that integrates NCBI E-Utilities, OpenAI’s GPT-4 and Zotero — to screen thousands of manuscripts and extract enzyme activities. Extensive validation revealed high precision and recall of GPT-4 in determining whether the abstract of a given paper indicates presence of a characterized enzyme activity in that paper. Provided the manuscript, FuncFetch extracted data such as species information, enzyme names, sequence identifiers, substrates and products, which were subjected to extensive quality analyses. Comparison of this workflow against a manually curated dataset of BAHD acyltransferase activities demonstrated a precision/recall of 0.86/0.64 in extracting substrates. We further deployed FuncFetch on nine large plant enzyme families. Screening 27,120 papers, FuncFetch retrieved 32,605 entries from 5547 selected papers. We also identified multiple extraction errors including incorrect associations, non-target enzymes, and hallucinations, which highlight the need for further manual curation. The BAHD activities were verified, resulting in a comprehensive functional fingerprint of this family and revealing that ∼70% of the experimentally characterized enzymes are uncurated in the public domain. FuncFetch represents an advance in biocuration and lays the groundwork for predicting functions of uncharacterized enzymes.

**Availability and Implementation:** Code and minimally-curated activities available at: https://github.com/moghelab/funcfetch and https://tools.moghelab.org/funczymedb

## Introduction

The last decade has witnessed a rapid rise in the number of high-quality sequenced plant genomes. However, most genes in these genomes are insufficiently annotated at the functional level. For example, there are 101 annotated BAHD acyltransferase enzyme-encoding genes in the cultivated tomato genome, out of which <15 have acceptor substrate classes associated with them (Kruse *et al*., 2022). While traditionally performed using sequence similarity searches across species using software such as PAINT (Gaudet *et al*., 2011) and TreeGrafter (H. Tang *et al*., 2019), in recent years, several methods employing artificial intelligence/machine learning (AI/ML) have also been developed for predicting gene function attributes such as Gene Ontology (GO) categories, Enzyme Commission (EC) numbers or enzyme-substrate information (Boadu *et al*., 2023; Pan *et al*., 2023; Kim *et al*., 2023). These algorithms typically utilize large public protein function databases such as UniProt, BRENDA and KEGG for their training data. Unfortunately, experimentally verified gene functions — mostly from a limited number of reference species — have been curated into these databases. Some model species such as *Escherichia coli* (Karp *et al*., 2023), *Saccharomyces cerevisiae* (Engel *et al*., 2022), *Schizosaccaromyces pombe* (Rutherford *et al*., 2024), *Drosophila melanogaster* (Antonazzo *et al*., 2020), *Arabidopsis thaliana* (Reiser *et al*., 2024), *Mus musculus* and *Homo sapiens* have strong curation communities, leading to prioritized deposition of experimentally characterized activities from these species into databases (de Crécy-lagard *et al*., 2022). However, despite a large corpus of experiments in other species, these characterized functions remain undeposited in function databases.

A major bottleneck for this deposition is biocuration (de Crécy-lagard *et al*., 2022). For example, of the >16 million entries in UniProt under Viridiplantae taxonomy class, only ∼43,000 have been curated in SwissProt (half of which are from Arabidopsis and rice), and only ∼12,000 entries have functions or catalytic activities defined based on literature evidence. This biocuration process involves — among a multitude of steps (Y. A. Tang *et al*., 2019) — searching for relevant papers, reading through them carefully, extracting relevant information into a structured database format, and validating the extraction. This is an invaluable but labor-intensive process. By one report, *“there are fewer than 100 full-time equivalents biocurators extracting gene-specific functional information from the literature into* ∼*40 public databases (functional/phenotypes/interactions/pathways) and fewer than 10% of these focusing on bacteria and plants”* (de Crécy-lagard *et al*., 2022). The small number of biocurators — especially in the plant domain — has resulted in a significant backlog of published manuscripts whose data is not available in structured databases.

In this context, large language models (LLMs) such as GPT, Gemini and Claude have emerged as promising tools for the automatic identification and extraction of scientific information from literature. General purpose models without domain-specific pretraining have been evaluated on a variety of tasks. For example, researchers have demonstrated that Claude 2 can extract structured information for medical evidence synthesis, highlighting its potential in therapeutic research (Gartlehner *et al*., 2024). A comprehensive assessment of GPT-3.5 revealed that while it falls short of state-of-the-art models for tasks such as named entity recognition and sentence similarity, it still nears human performance on the PubMedQA evaluation (Chen *et al*., 2023).

Additional work has evaluated the performance of LLMs that are fine-tuned or pretrained on domain-specific text. One study demonstrated that an older model, GPT-3, could perform similarly or outperform dedicated models on a variety of evaluations including questions on molecule properties and chemical reaction yields (Jablonka *et al*., 2024). Similarly, LLMs enhanced with explicit chemistry knowledge outperformed others in chemistry-related tasks, including organic synthesis and the discovery of novel chromophores (Bran *et al*., 2023). GPT-4 and BERT-based models have also shown strong performance in establishing protein-protein interactions despite the lack of domain specific text training for GPT-4 (Rehana *et al*., 2023). A comparative study of ML models indicated that a fine-tuned PubMedBERT based model was superior in identifying gene-disease relationships when compared to rule-based systems and T5-based models (Milošević and Thielemann, 2023). This growing body of knowledge establishes LLMs as valid tools for interacting and gleaning valuable data from corpuses of biomedical text.

While ongoing work continues in development of models, evaluation frameworks, and benchmark performance, there are opportunities to deploy readily-accessible LLMs for useful information retrieval tasks. One workflow demonstrated success in phylochemical mapping of specific natural products to plant phylogenies (Busta *et al*., 2024). The PATHAK method successfully elucidated gene function relationships in *Arabidopsis thaliana* from ∼5000 journal articles (Kumar and Mukhtar, 2024). These efforts suggest that current generation of LLMs already have sufficient utility for information retrieval applications. Despite these successes, to our knowledge, the large corpus of literature on plant enzyme function and enzyme-substrate relationships going back four decades has still not been parsed. Furthermore, the challenges associated with the quality of such extractions have not been adequately explored.

In this research, we evaluated the capability of GPT-4 for extracting enzyme-substrate information and related metadata from published papers. We developed FuncFetch, a multi-step workflow to generate such an enzyme-substrate dataset. We demonstrated the utility of this workflow by controlled testing and iterative development and compared its performance relative to a high-quality, manually curated enzyme-substrate database of a large enzyme family. FuncFetch is adaptable to other gene families with a few prompt changes. We demonstrate this extensibility by producing a first-pass curation dataset of >30,000 characterized enzyme-substrate interactions from nine plant enzyme families, which together constitute ∼5% of the diploid plant genomes. Further manual curation of this dataset can enable better annotation of thousands of enzyme-encoding genes across the plant kingdom.

## Methods

### Dependencies

The multi-step FuncFetch workflow **(Fig. 1A)** was developed using Python v3.10.13 and requires installation of specific additional packages **(Supp. File 1A)**. Additionally, it requires OpenAI and NCBI accounts and Application Programming Interface (API) keys. All configuration files and code can be found on our GitHub page (see Data Accessibility).

**Figure 1:**
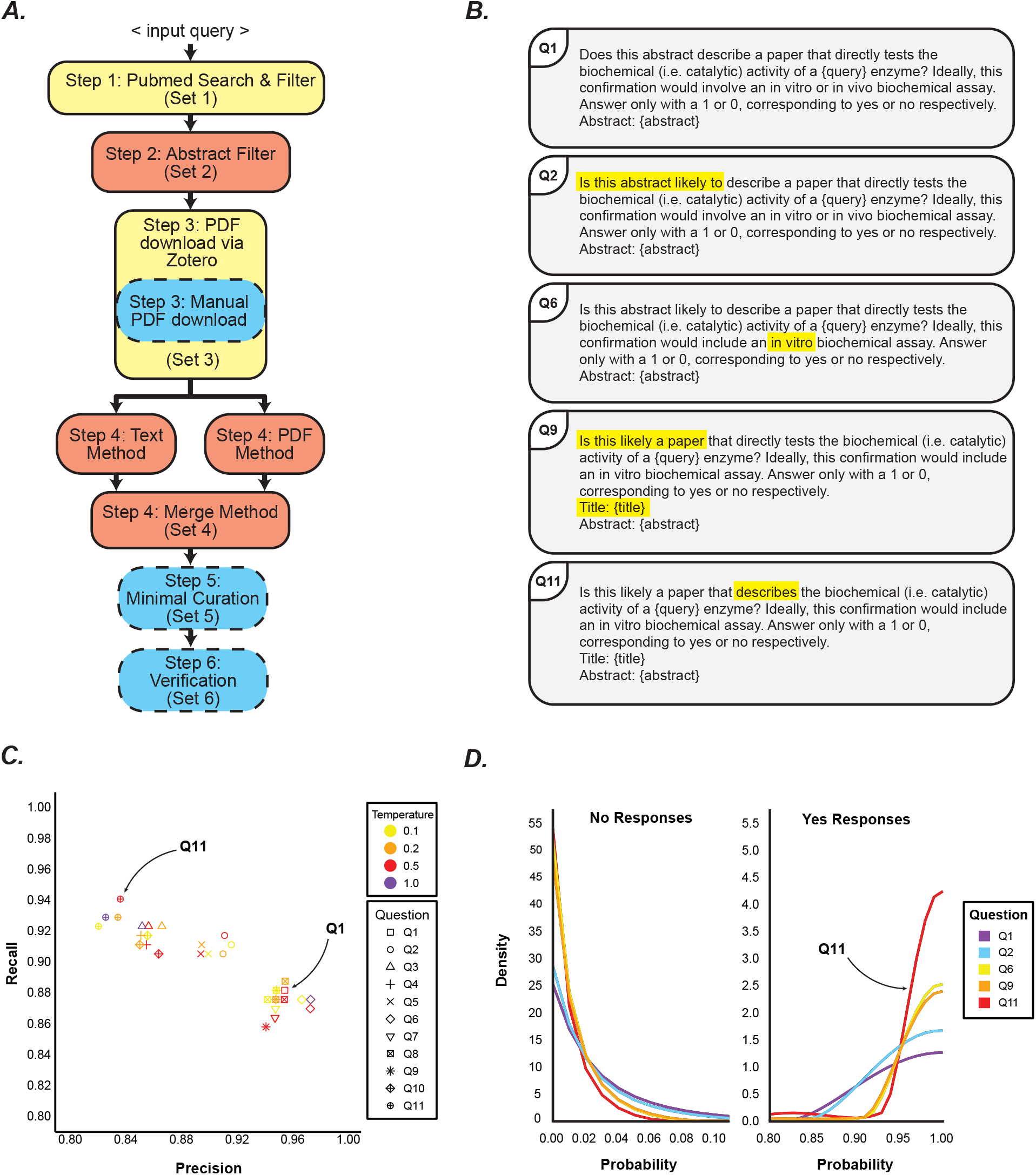
Overview of pipeline and performance assessment. (A) Pipeline overview. Boxes in yellow show steps not requiring GPT-4, those in salmon require GPT-4, and those in blue are executed manually or using local specialized Python scripts. (B) Representative questions for Step 2. Yellow highlight indicates altered text relative to the above question. Full list of questions available in Supplementary materials. (C) Performance assessment of query engineering. Q11 question with the highest recall at temperature 0.5 and was chosen for downstream steps. Q11 provided more flexibility for the model in predicting target papers. (D) Density estimation of probability value distribution indicating increased model confidence in “yes” (“1”) responses. Boundary-corrected kernel density estimates computed using biweight kernel and simple boundary correction method (fbckden function from evmix R package). Note that the Y-axes for no and yes response plots have different scales.

### FuncFetch overview

This workflow comprises of five steps as follows:

- **Step 1:** With API calls to NCBI E-Utilities (Sayers, 2018), PubMed was queried using a user-supplied list of journals and keywords **(Supp. File 1B-D)**. The keywords must first be decided by the user based on domain-specific knowledge using the PubMed web interface. Given our focus on plants, the journal list included 199 plant, biochemistry and general interest journals. By default, all review articles and all only-review journals were excluded from further consideration but the users have the ability to include these article types. The articles were filtered by a list of 156 keywords related to biochemistry. Given limitations of the NCBI efetch pipeline, if there were >10,000 hits, multiple queries were made with publication-year limits, enabling download of all hits systematically. These settings can be adjusted using a configuration file.
- **Step 2:** Abstracts of articles selected from Step 1 hits (Set 1) were passed on to GPT-4 Turbo (gpt-4-turbo-2024-04-09), which was given the abstract text, title and a family-specific query. The model returned a yes or no response of whether the paper likely described the biochemical activity for the given enzymatic activity. For papers classified as yes, the software output a set of Digital Object Identifiers (DOIs) and other citation information in RIS format (Set 2).
- **Step 3:** This step was performed in a semi-automatic fashion. The RIS file was imported into the open-source program Zotero 6.0.32 or v7 beta, which automatically downloaded PDFs of the manuscripts locally. Manual download was needed for some articles.
- **Step 4:** The PDFs in the Zotero storage folder were passed to GPT-4 directly as well as after text extraction using pdfminer.six v.20231228. Specific prompts **(Supp. File 4)** were sent to GPT-4 using API calls to extract the enzyme-substrate pairs as well as other annotation information such as enzyme common name, species name, GenBank/UniProt ID if any, and product information. The API calls were made with an associated vector store containing a single PDF file for the PDF method. The text method sent extracted text to the model as a string in the API request. Information parsed from both approaches was merged via another GPT-4 API call. This API call was made using a third prompt along with the string outputs of both previous methods. The prompts instruct the model to output data in JSON format, which was then converted to a tab-delimited file. We evaluated both GPT-4 Turbo and GPT-4o models (gpt-4-turbo-2024-04-09 and gpt-4o-2024-05-13, respectively). This step produced a set of extracted enzyme activities (Set 4, uncurated set).
- **Step 5:** Manual verification of Set 4 activities is essential because Set 4 can contain plant enzymes tested in bacteria/yeast via heterologous expression, actual non-plant enzymes that need to be removed, or enzyme activities from a different family. We did not test the ability of GPT-4 to identify these complex scenarios. In this step, for each row of extracted Set 4 activities, if a species name was available, its taxonomic information namely the family and kingdom were attached using a custom Python script. Using Microsoft Excel and custom Python scripts, subsequent curation involved selecting enzymes with species information, removing activities from non-target families, adding UniProt and GenBank IDs based on extracted ID information, and prioritizing enzymes with available gene IDs for further curation (Set 5, minimally curated set).
- **Step 6:** This step involved reading each prioritized paper manually and verifying extracted data in each column, resulting in Set 6, a Verified Set. In this research, Step 6 was performed only for BAHD acyltransferase enzymes.

### Validation of Step 2

A test dataset of 347 abstracts was manually constructed to equally represent papers with reported enzyme activities (positives) and no reported enzyme activities (negatives) **(Supp. File 2A**,**B)**. Positives were sourced from UniProt citations for catalytic activity annotations, with additional papers identified through manual searches to ensure equal representation across five enzyme families. We included these enzyme families — BAHD acyltransferases, cytochrome P450s, O-methyltransferases, UDP-glycosyltransferases, peroxidases — to reduce bias and overfitting of prompts to any one family or activity. The abstract and title of each paper were assessed by an API call to GPT-4 along with a predetermined prompt embedded with the enzyme family name **(Supp. File 2B)**. The model parameters were: *seed = 1, logit_bias = {‘15’:100, ‘16’:100}, max_tokens = 1*. The seed was set to increase reproducibility of model outputs during validation. The logit_bias and max_tokens parameters limit the model’s output options to a single token response of “1” or “0”, corresponding to yes or no respectively. Model temperatures were tested at 0.1, 0.2, and 0.5 based on OpenAI’s recommendation for tasks that require consistent outputs (OpenAI Docs, 2024). Based on the output of the model, each paper was established as true positive, false positive, true negative, or false negative enabling calculation of precision, recall, and F1 scores for each prompt across the entire dataset. Thirteen unique prompts were developed throughout testing **(Supp. File 2B)**. This began with an initial prompt developed by interacting with ChatGPT and asking it to extract biochemical information from pdfs of journal articles. Prompts were developed by reading through the output of previous prompts and making hypotheses about the most relevant language in the prompt that influenced the false positives/negatives. The system content of the chat completion model was also iterated upon, but seemed to make little difference in performance, so evaluation of this parameter was stopped. Whenever a new prompt resulted in a better F1 or Recall score than a previous attempt, that prompt would then be iterated upon via the same process.

### Validation of Step 4

A test set of 10 papers was established to evaluate the output and performance of FuncFetch Step 4 **(Supp. File 3A, Supp. File 4)**. This test set was made up of 5 papers each from two enzyme families in the positive paper set detailed in the previous section (BAHD acyltransferases [BAHDs] and UDP glycosyltransferases [UGTs]), containing 71 and 38 BAHD and UGT activities from 16 and 18 enzymes, respectively. These ten papers were selected to represent the diversity of BAHD activities and contexts in which they were reported (e.g. single enzyme discoveries, complete biosynthetic pathway studies, heterologous expression). The UGT papers were used as the negative test with the query set as “BAHD acyltransferase” in the Step 4 configuration. This ensured that there were enzyme activities present in the paper that could be extracted successfully, but that would not be of the correct enzyme family. We included this set of papers as the negative test set to evaluate the model’s ability to discriminate enzyme families and exclude irrelevant activities. Step 4 was run on these 10 papers and the outputs of text, pdf, and merge extraction techniques were graded for precision and recall.

All extraction methods output multiple activities in the same line of tabular output, but especially the merge method. Given this model behavior, evaluating each line of output underestimates the number of correct activity relationships. Therefore, we developed a manual procedure that splits a line of output into multiple entries. A grading procedure was developed to score entries of Step 4 output as correct, unknown correct, incorrect, or incomplete **(Supp. File 5B**,**C)**. These procedures also detail enzyme, substrate, and product naming standards. Graded entry counts were used to calculate evaluation metrics.

### Assessment of workflow using curated BAHD acyltransferase database

After validation of the entire pipeline, FuncFetch was used to extract information about the BAHD family, resulting in Set 4 enzyme-substrate association entries. The tabular output of Step 4 was evaluated against a high-quality dataset (HQD) of BAHD activities **(Supp. File. 5D)** from 129 papers updated from our previously published study (Kruse *et al*., 2022). Every entry in this dataset was manually reviewed at least twice. This HQD represents the best comparison to the performance of FuncFetch Step 4 because there are no comparable datasets available and because FuncFetch aims to complement family-wide manual curation efforts. For each paper appearing in the evaluation set and the HQD, the Step 4 output was manually graded and compared to the HQD according to the same grading procedure described in the previous section.

### Semi-automatic curation to generate the minimally curated set

Set 4 entries were processed in a multi-step fashion. First, we flagged and manually reviewed a small subset of entries in the tabular output for which the activity extractions came only from the PDF method or text method or none. Second, we identified records of plant proteins either directly ascribed to plants or tested in bacterial or fungal systems. This was accomplished by tagging each row of the Set 4 result with the Family and Kingdom name using NCBI Taxonomy hierarchy. Third, we identified records of the targeted enzyme family using the extracted enzyme names, abbreviations and if needed, reading the papers. Non-specific extractions (e.g. cytochrome P450s or hydroxylases or acyltransferases when the target family was methyltransferases) were removed. Fourth, we used sequence IDs and enzyme names to match entries with UniProt accessions. Finally, entries with sequence ID information available were prioritized, given the ability to identify their enzyme families with certainty using protein sequence information. These entries with sequence ID information constituted the Minimally Curated Set (Set 5). Only BAHD acyltransferases were processed to Verified Set (Set 6), which involved a detailed reading of the associated literature and adding/deleting activities as needed **(Supp. File 6)**. We note that there are substantial similarities in activities extracted for the CYP450 and dioxygenase/oxygenase queries, given significant overlap and promiscuity in functions of these families.

### Development of a web resource and data availability

All scripts used in this work are available on GitHub (https://github.com/moghelab/funcfetch) along with the demo files, while relevant output files were deposited on GitHub and the project website (https://tools.moghelab.org/funczymedb/). For developing the website, rapid prototyping of page layouts and functionality was initially performed in Webflow (2024) before development was moved and completed in VSCode v1.87. Node.js v20.12.0 was used to generate and launch the pages as a web app. This Node.js app is dependent only on the Express module v4.18.3. The website was run in a Docker v20.10.17 container built from a Node.js base image: *node:16*.*17*.*0-bullseye-slim*. The base image acts as the parent of the image, and has heritable capabilities.

## Results and Discussion

### FuncFetch enables first-pass extraction of enzyme-substrate interactions from manuscripts

FuncFetch Step 1 queries the NCBI PubMed database using the user-provided search term and identifies manuscripts containing or associated with the query as keywords or MeSH terms. Preliminary assessments revealed that many of these hits are not relevant e.g. a query for BAHD acyltransferase also retrieved articles on Bone-Anchored Hearing Devices, or on topics related to plant development. Hence, we instituted filters based on journals and keywords **(Supp. File 1B**,**C)**. Considering hundreds of irrelevant manuscripts and manuscripts with only mention of the enzyme but no detailed *in vitro* investigation still passed these filters, we explored whether GPT-4 can predict whether a given paper might describe an enzyme assay for the family of interest, based on the title and abstract text. We manually generated a corpus of 347 manuscripts belonging to five enzyme families to test GPT-4’s performance.

Our first prompt **(Q1; Fig. 1B)** resulted in recall, precision, and F1 scores of 0.88, 0.95 and 0.91, respectively, at temperature 0.5 **(Fig. 1C)**, indicating an acceptable level of performance. To optimize it further, we sequentially iterated through 11 questions and 2 system inputs in 13 combinations **(Fig. 1B; Supp. File 2B**,**C)** revealing an expected tradeoff between precision and recall. We selected Q11 as it had the highest recall (0.94), enabling us to cast a wide net for the next step. We also saw that introducing “likely” (Q1 to Q2) and “describes” rather than “directly tests” (Q9 to Q11) resulted in increased papers classified as relevant, from 156 to 170 and 154 to 190 respectively. The density of “yes” responses increased for prompts with more inclusive language **(Fig. 1D)**, highlighting the relationship between the model’s confidence in assertions and the specificity or flexibility of prompt language.

Having demonstrated the validity of the Abstract Filter, the next step required parsing the entire manuscript. We assessed that the best solution to accessing the PDFs is to export a citation exchange RIS file and download papers automatically using the Zotero reference manager. This step was successful for a majority of the cases — using Zotero v6, 74.8% of BAHD papers could be automatically downloaded. Remaining papers were downloaded mostly one-by-one and attached to Zotero records manually.

Initially we tested two approaches for accurate retrieval of enzyme activities from downloaded manuscripts **(Supp. File 4)**: 1) directly feeding in a PDF to GPT-4 (Approach 1), and 2) embedding PDF data in a text format, followed by feeding the text to GPT-4 (Approach 2). Comparative testing of these two approaches revealed that Approach 1 had better recall than Approach 2, with precision being better for the latter **(Supp. File 3B)**. Combining the outputs of both approaches again using an LLM instruction (Approach 3; “merge method”) resulted in improved performance **(Supp. File 3B; Supp. File 4)**.

### Performance of the FuncFetch pipeline on the BAHD acyltransferase dataset

We next implemented the entire pipeline on the BAHD acyltransferase enzyme family dataset. We previously compiled a dataset of 567 activities of 164 BAHDs from 75 species (Kruse *et al*., 2022), which we updated in this study. The final HQD comprised 1114 activities of 209 BAHDs from 87 species **(Supp. File 5D)**. We estimated precision and recall of the FuncFetch pipeline by comparing its output with the HQD. The total PubMed articles returned by Step 1 were 1038 with elink and 461 without elink **(Fig. 2D)**. Of the 129 papers in the HQD, 113 (87.6%) and 78 (60.5%) were retrieved, with and without elink respectively. Of these, Step 2 classified 101 and 76 as relevant (Recall: 0.89 and 0.97). Seventy additional papers with 137 novel characterized BAHDs were found by this workflow and missed by our manual search.

**Figure 2:**
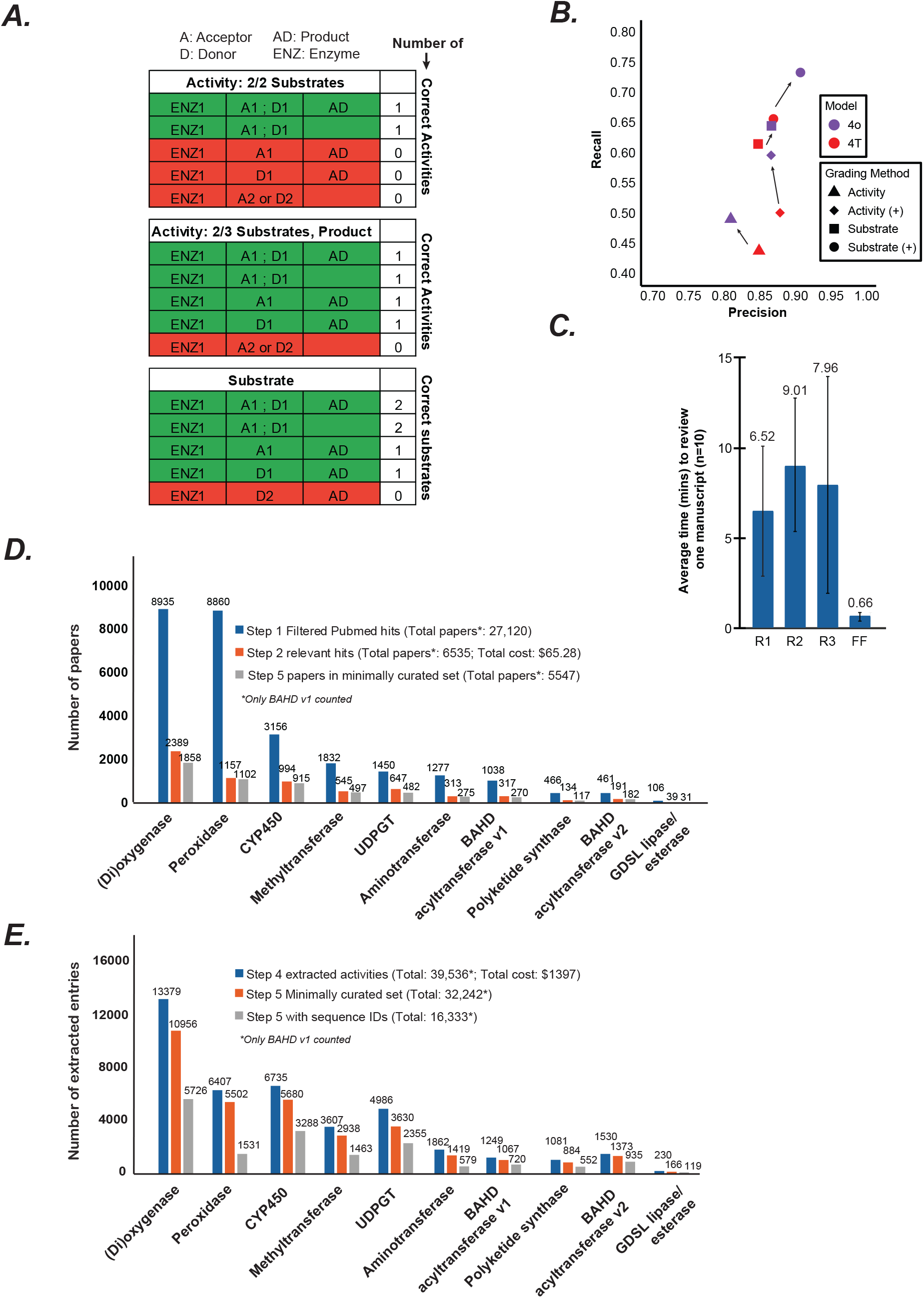
FuncFetch results. (A) Schema for Step 4 grading procedure. Red and green colors depict wrong and correct entries, respectively. (B) Precision-Recall of Step 4 output. *Activities* are entries correct on 2/3 compounds present in a reaction. *Substrates* are counted as correct or incorrect independently from activity grading. (+) denotes entries regraded with unknown correct substrates or product counts added to the high-quality dataset (HQD). (C) Average time taken by three human reviewers [R1, R2, R3] and FuncFetch Step 4 [FF] to extract information from 1 manuscript. (D) Number of papers obtained at different steps for each family. The keywords used for initial extraction are described in Supplementary File 1. Costs were calculated by noting the difference between API credits used before and after each code run. (E) Number of extracted and minimally curated enzyme-substrate entries. Entries with sequence IDs can further assist in verifying the domain family of the extracted enzyme. Significant overlap was seen between entries extracted for dioxygenases and CYP450s.

From the 76 papers passed on to Step 4, we estimated reaction-level and substrate-level performance metrics using the HQD as the comparison. The precision/recall was 0.81/0.49 and 0.87/0.65 respectively **(Fig. 2B)**, indicating that while GPT-4 was correctly able to extract the substrates most of the time, it faced challenges linking the substrates and/or substrate-product pairs together. We also found that the performance of both reaction and substrate-level extraction increased with a model update from GPT-4T to 4o **(Fig. 2B)**.

The most common cause of an incorrect entry in the output were cases of incorrect enzyme families. Of 65 false positive entries, 20 (30.8%) were BAHD family enzymes with incorrect activities not supported by the text. The other 45 (69.2%) incorrect entries were enzymes of non-target families. These other families included acetyltransferases, decarboxylases, carboxylases, UDP-glycosyltransferases, O-methyltransferases, ligases, and various synthases. Papers resulting in these entries discuss entire biosynthetic pathways or otherwise report on a variety of enzymes and reactions. Despite some confusion on enzyme families, the model displayed an ability to understand domain-specific nomenclature. For example, the model parsed and associated CoA thioester naming conventions (e.g. aiC4, iC5, aiC5) with products and other substrates effectively **(Supp. File 5J)**.

In our evaluation, 191 (24.0%) of the 795 entries were marked as incomplete. These generally tended to be incomplete donor substrates rather than acceptors. Of 96 incomplete entries with one substrate, 74 (77.1%) were missing a donor and 22 (22.9%) were missing an acceptor. These omitted data may result from additional attention paid to acceptors due to similar naming of acceptors and products, and the separation of donor information in the text from discussion of acceptor substrates and products.

Of the relevant entries, ∼58% of entries with extracted IDs/names could be associated with at least one UniProt entry. These entries can be considered priority entries for more detailed verification **(Table 1)**. We sampled 10 papers with verified activities that were missing sequence IDs to understand why these IDs were omitted. In four papers, enzymes lacked IDs, though sequences were available in supplementary files or external references. Three papers described enzymes not mapped to genes, so no sequence information was available. In the final three, IDs were present but missed during retrieval. These findings highlight areas where our ID retrieval could be improved, though some IDs may remain unavailable with our current method.

**Table 1:** Extracted entries that could be mapped to one or more UniProt IDs.

| Enzyme family name | Total extracted entries | Number of entries with ID hits | % with ID hits | Number of entries without ID hits |
| --- | --- | --- | --- | --- |
| GDSL lipase | 166 | 97 | 58.4 | 69 |
| UDPGT | 3630 | 2089 | 57.5 | 1541 |
| BAHD acyltransferase | 1429 | 841 | 58.6 | 588 |
| Peroxidase | 5502 | 2820 | 51.3 | 2682 |
| OMT | 2938 | 1401 | 47.7 | 1537 |
| CYP450 | 5680 | 3558 | 62.6 | 2122 |
| Aminotransferase | 1419 | 740 | 52.1 | 679 |
| Dioxygenase | 10957 | 6668 | 60.9 | 4289 |
| PKS | 884 | 624 | 70.6 | 260 |
| **TOTAL** | **32605** | **18838** | **57.8** | **13767** |

Out of 795 Step 4 output entries, we found 237 correct entries that were not found in the manual curation process **(Supp. File 5J)**. These 237 entries could be associated with 29 UniProt accessions. These data include activities inferred from enzyme names, references to previous assays performed in the references section of the manuscript, and activities assayed but entirely missed by the manual review process. When we evaluated Step 4 output including these entries, precision/recall increased at the reaction-level and substrate-level, to 0.87/0.60 and 0.90/0.73, respectively **(Fig. 2A,B; Supp. File 5A)**. This finding highlights the value of a complementary LLM screening of manuscripts along with manual curation, as a human reviewer may overlook information the model detects.

We also evaluated the proportion of our retrieved entries that are already present and curated in UniProt i.e. in the SwissProt and RHEA databases. This provided a sense of the “curation gap” that is addressable by FuncFetch. From the 841 verified BAHD entries with extracted UniProt IDs, we found 187 unique IDs **(Table 1)**. Of these unique IDs, 49 (26.2%) were reviewed (Swiss-Prot) and 44 (23.5%) had catalytic activity (RHEA) annotations **(Supp. File 6A**,**B)**. We also missed 20 BAHDs already catalogued in SwissProt and RHEA **(Supp. File 6B-D)**. Considering these numbers, we estimate that at least 69.1% of the Verified Set lacks catalytic activity annotations. This represents the current estimated curation gap for this enzyme family, which is likely an underestimate given 588 extracted entries could not be automatically assigned a UniProt ID and were not included in this calculation. This analysis demonstrated that our workflow is capable of finding enzyme activities that are already well-represented in public databases. However, most entries retrieved by this workflow remain uncurated in the public domain, validating the utility of our approach.

### First-pass automatic curation of thousands of manuscripts for eight plant enzyme families

To extend our analysis further and generate a valuable resource for the plant biochemistry community, we used FuncFetch to identify relevant papers for eight additional enzyme families/activities that constitute ∼4-5% of the gene content in diploid plant genomes **(Fig. 2C-E)**. This process was an order of magnitude faster than manual review **(Fig. 2C; Supp. File 7)**. At a cost of ∼$65, we screened >27,000 paper abstracts and identified a conservative set of 5547 manuscripts that may contain enzyme-substrate information **(Fig. 2D)**. Performance of Steps 3-5 produced a Minimally Curated Set for each family **(Fig. 2E)**. Overall, across all 9 enzyme family activities, 32,605 entries were extracted and minimally curated from 5547 papers, which will need to be verified further. Detailed Step 6 curation was performed only for BAHD acyltransferases **(Supp. File 6)**.

Curation performed in Step 5 to produce this Minimally Curated Set revealed several issues with LLM-extracted data that needed to be carefully assessed manually. First, we found that 10-30% entries for each family included non-target enzymes from other families or non-plant enzymes. Without manual curation, it is not straightforward to identify plant enzymes, since some papers may contain plant enzymes heterologously expressed in bacteria or yeast. In several cases, the model listed the host species name (e.g. *Escherichia coli, Saccharomyces cerevisiae*) rather than the name of the original plant species. Second, if multiple enzymes are characterized in the same study, multiple enzyme entries could be extracted, and need to be filtered out. Third, we observed a consistent gap in retrieved enzyme sequence information across all families **(Table 1)**. On average, ∼42% of entries have no associated sequence identifier after we map them to UniProt accessions based on enzyme names. This presents a roadblock to eventual manual curation.

In addition to extraction issues, we detected several instances of hallucinations **(Supp. File 8)**. For plant papers and enzymes, the model listed human, mouse or rat as species and listed their enzymes and substrates despite these species never being mentioned in the paper. While there were 0-1% hallucinations detected for plant-enriched families such as BAHDs, PKSs, GDSLs and UDPGTs, the proportion was between 3-7% for ubiquitously present families such as peroxidases, CYP450s and (di)oxygenases. The actual number of hallucinations of this type could be higher but is irrelevant given they would be filtered out using the taxonomic filter. The tabular organization of the data in the Step 4 output and association of paper titles with species and kingdom names in Step 5 allows for efficient detection and removal of many such hallucinations. Nonetheless, detailed curation needs to be performed for all entries before their release on protein function databases. Minimally Curated Sets for all enzyme families can be accessed and downloaded from the project website and GitHub.

## Conclusions

Since the introduction of transformers in 2017 (Vaswani *et al*., 2017, 2023), the field of LLMs has grown by leaps and bounds as witnessed by the numerous commercial LLMs existing today including GPT, Llama, Mistral, Gemini, and Claude. Biological research has made significant use of these LLMs just in the last two years in fields as diverse as predictions of protein-protein interactions (Jin *et al*., 2024), single-cell multiomics clusters (Cui *et al*., 2024), summarizing literature for non-coding RNAs (Green *et al*., 2024), predicting disease specific knowledge graphs (Li *et al*., 2024), and in eco-evolutionary research (Gougherty and Clipp, 2024). Here, we explore the use of the GPT-4 LLMs in extracting enzyme-substrate interactions and related metadata, highlight the limitations of this application, and generate a resource of such interactions for community annotation and use. Such an application has been explored recently e.g. with ChatGPT and Bard/Gemini in a microbiological context (Caspi and Karp, 2024) and for general entity-relation pairing in PubTator3 (Wei *et al*., 2024). Kumar and Mukhtar developed a method named PATHAK for associating genes with biological process GO terms (Kumar and Mukhtar, 2024). However, this is a dynamic field and a state-of-the-art application, as highlighted by the fact that many of the cited approaches were still preprints at the time of this writing. To our knowledge, at the time of writing, no existing workflow exists for extracting plant enzyme activities from biochemistry literature. In addition to describing a workflow, this work also makes a list of hundreds of relevant papers and thousands of partly curated activities available to the scientific community for community curation/analyses. Given over two-thirds of the experimentally characterized enzymes are still uncurated in the public domain, development of workflows such as FuncFetch is a need of the hour.

The FuncFetch workflow integrates three platforms — NCBI E-Utilities, OpenAI GPT-4 and Zotero — enabling screening of thousands of published papers and extraction of information from a select few. This workflow can be flexibly configured as needed for other purposes. We expect that the overall sequence of steps will remain constant, although the questions as well as the precision/recall for different tasks may change based on task complexity. For example, identifying chemical and enzyme names — both of which have a unique, defined vocabulary — may be a simpler task for an LLM than extracting mutant phenotype data, which could range from change in cellular morphology or membrane fluidity to change in growth rate or interaction with pests/pathogens. Therefore, query engineering may need to be conducted for every untested application of FuncFetch. Over the course of this research, we also found that Step 4 performance and especially cost-effectiveness significantly improved with GPT-4 model updates **(Fig. 2B)**. We note that FuncFetch is flexible enough to accommodate future model upgrades. Assuming that models will keep improving over time, the cost-effectiveness and performance of the FuncFetch pipeline can be expected to improve.

The advantage of the stepwise approach is evident. It took ∼14 minutes to download >27,000 paper metadata from PubMed, ∼$65 to screen these abstracts overnight using GPT-4 batch submission and ∼$1400 to extract ∼40,000 entries. The initial screening step significantly reduced the cost of data extraction from PDFs, and the pre-selection of papers mitigates the chances of citation hallucinations. One major bottleneck in our pipeline is downloading PDFs of screened papers using Zotero (Step 3). This is not the best option; however, it is the most optimal option that circumvents manually downloading thousands of papers while still retaining comprehensiveness. We expect that individual institutions’ access to publisher libraries may affect the outcome of this step. For Step 4, we compared the cost of manual curation vs. GPT-4 extraction from papers. The differences in cost and time savings were drastic **(Fig. 2C; Supp. File 7)**. We believe these are gross underestimates since the effect of human attention span and engagement over long periods of time are not accounted for here. Nonetheless, given the quality issues of GPT-4 extraction highlighted above, manual validation of extracted activities is still necessary.

We decided to use the GPT-4 LLM for this analysis given its widespread use, however, other LLMs listed above may offer better performance. In validating Step 4 methods we found some output variation between identical API calls, despite the inclusion of a seed parameter. These variations generally consisted of recombination of substrates, products, and enzymes across different entries. It may be worthwhile to explore these variations and leverage model information on probable alternative token responses to extend the breadth of extracted information. Furthermore, we observed that GPT-4o performed better than GPT-4T **(Fig. 2B)**. The choice of the LLM model and parameter changes may therefore significantly affect curation results. Recently, EnzChemRED emerged as a chemistry relation extraction dataset comprised of curated PubMed abstracts. Finetuning on such a dataset appears to improve model performance on relation extraction metrics (Lai *et al*., 2024). Taken as a whole, optimization of finetuning, model choice, and parameters will likely improve performance on similar information retrieval tasks.

Despite our initial concern, we found zero instances of hallucinations (Jin *et al*., 2023) in our full curation of BAHD activities, which we define here as making up substrates and products that are not mentioned in the paper. While the LLM made incorrect associations between substrates and enzymes, either from the main text or the references, completely un-mentioned substrates were not observed in Set 5 entries for BAHDs and PKSs. Nonetheless, for more ubiquitous families, dozens to hundreds of instances were detected. Hallucinations are known to occur when using LLMs, especially public-facing versions such as ChatGPT and Bard (Caspi and Karp, 2024). Our results suggest that in some cases, the model outputs information from the training vector space **(Supp. File 8)**. Such hallucinations tended to be highly repetitive and near identical, listed biomedical species instead of plant, bacteria or yeast, or had human enzyme names when the paper title clearly indicated investigation of a plant enzyme. This finding also raises a potential concern for chemistry or protein-function specific LLMs built on foundation models — the taxonomic breadth of the training data used may significantly affect their performance.

In FuncFetch, we employed several measures that potentially mitigate hallucinations. Firstly, our questions were highly specific in the enzymatic function as well as the taxonomic space. We also focused on single enzyme families, significantly constrained the outputs to 1/0 (Step 2) or a controlled JSON vocabulary (Step 4), and set low temperatures for retrieval (0.5 instead of default 1 on ChatGPT) thereby reducing model “creativity”. For each query in Step 4, we also initiated a new API call removing all memory of previous uploaded papers/text. Third, we employed extensive semi-automatic curation steps (Step 5) that lead to detection of obvious instances of such hallucinations. Finally, we matched the extracted species name with the species name of the sequence identifier obtained via NCBI Taxonomy, enabling us to select higher-confidence entries rapidly (Step 5). Nonetheless, it is possible that full curation of the Set 5 activities of the non-BAHD enzyme families may reveal additional instances of hallucinations. Future development of the FuncFetch workflow may require integration of techniques such as Retrieval Augmented Generation (RAG) (Béchard and Ayala, 2024; Wei *et al*., 2024), Knowledge Retrieval (Varshney *et al*., 2023) or LLM-finetuning for genomics-focused tasks. Mitigating hallucinations in a high-throughput manner is a dynamically evolving field at the time of this writing (Tonmoy *et al*., 2024).

ML has been used for predicting genes and gene functions for over two decades (Mahood *et al*., 2020). Over the last few years, however, the advent of AlphaFold and AI innovations have significantly revolutionized this field. For example, Google’s ProtNLM recently provided functional descriptors for 50 million uncharacterized proteins in the UniProt database (Hatch, 2022). ESP — a generalized machine learning model — was found to predict enzyme-substrate pairs with >91% accuracy for the tested datasets (Kroll *et al*., 2023). In-house transformers have also been used to predict enzyme-substrate promiscuity — AlphaFold’s EvoFormer model was used to predict the strength of protein-ligand interactions, with additions from orthogonal cheminformatic techniques (Xing *et al*., 2024). While these innovations are useful in covering the vast, unannotated protein space, our previous research on BAHDs demonstrates that simply cataloguing activities of published, well-characterized BAHDs increased the proportion of uncharacterized enzymes annotated with their putative substrate classes from ∼10% to 40-45% (Kruse *et al*., 2022). Furthermore, such a catalog allows more rational selection of substrates for downstream validation experiments as well as improves the training data for generation of more unbiased models. The FuncFetch workflow and database are steps in that direction.

## Acknowledgements

We thank Drs. Qi Sun and Jaroslaw Pillardy at the Cornell BioHPC core and Vishwa Jyoti Baruah for their assistance in implementing the pipeline and the website Docker container. We thank Venus Kajangu for assisting with verifying GenBank sequence IDs in Set 6 for BAHDs. We also thank the reviewers for helpful comments that helped improve this manuscript.

## Author Contributions

This research was conceived by NS and GM. NS, CM and GM wrote code and performed all bioinformatic analyses. NS, XY and GM validated results and performed curation. NS and GM wrote the manuscript. All authors reviewed the manuscript.

## Supplementary Data

**Supplementary File 1:** Description of packages and filters used for FuncFetch workflow. (1A) Version information for all software used to develop and test FuncFetch pipeline. (1B) PubMed indexed journals included by FuncFetch when running Step 1 with Journal Filter. (1C) PubMed indexed keywords included by FuncFetch when running Step 1 with keyword filter. Keywords marked with “fil” in the second column were not used for most families but are kept in the list in case needed. (1D) Queries used for PubMed searches.

**Supplementary File 2:** Details of Step 2 testing & results. (2A) PubMed IDs and titles of 347 papers included in evaluation of Step 2, and whether in positive or negative test group. (2B) Text of all prompts and system input evaluated for Step 2. (2C) Metrics of every prompt and system tested across temperatures (Descending F1 score order).

**Supplementary File 3:** Details of Step 4 validation. (3A) List of 10 papers included in Step 4 method validation. Papers in the positive set have BAHD acyltransferase activities. Papers in the negative set have UDP-glycosyltransferase activities, but the model was asked to extract BAHD acyltransferase activities. (3B) Metrics for each tested method using the 4T or 4o models.

**Supplementary File 4:** Details of Step 4 prompts and code examples.

**Supplementary File 5:** Details of Step 4 Evaluation. (5A) Analysis and results of Step 4 of FuncFetch workflow. Counts of unknown correct (dark blue), known correct (green), incorrect (red), and incomplete (yellow) entries for each evaluation method. Additionally, counts for entries that differ in grading between evaluation methods denoted in a separate section as dark blue, light blue, and orange. Recall and precision scores for each evaluation method, and formulas used to calculate each. (5B) Protocols for grading Step 4 output entries. This includes definitions and examples of what constitutes correct and incorrect entries and substrates/products. (5C) Protocol for splitting Step 4 output entries. This process splits an entry with multiple combinations of substrates (activities) to align Step 4 output with the structure of the high-quality dataset [HQD]. (5D) HQD of BAHD acyltransferase activities. (5E) Tabular output file of Step 4 (merge method) run with model gpt-4-turbo-2024-04-09 (4T). (5F) Tabular output file of Step 4 (merge method) run with model gpt-4o-2024-05-13 (4o). (5G) HQD entries from papers that appear in output of 4T Step 4. (5H) HQD entries from papers that appear in output of 4o Step 4. (5I) 4T Step 4 output entries from papers that appear in the HQD. (5J) 4o Step 4 output entries from papers that appear in the HQD. (5K) 4T Step 4 output substrate grading. (5L) 4o Step 4 output substrate grading. **Supplementary File 6:** Verified Set of BAHD acyltransferase activities. (A) The Manual Check column differentiates entries where the species of the FuncFetch extracted ID is the same as the one extracted through NCBI Taxonomy using the enzyme names by our inhouse Python script. If the tag is “Same Species”, the UniProtID column names apply. If the tag is “Different_species” or “Multiple_species”, then none of the NCBI Taxonomy IDs extracted are correct, and we consider them incorrect mappings to be dealt with manually. If the tag is “No_ID”, then No ID could be extracted. In total, out of the 1429 entries, IDs of 841 entries could be extracted, only 53 of which are incorrectly mapped after Manual_Check. (B) Analysis of IDs for determining Curation gap. (C) All SwissProt BAHDs (D) Overlaps between different sets.

**Supplementary File 7:** Time comparison of human readers and Step 4. (7A) Time totals, analysis, and methods. Human readers recorded the time it took them to read and extract enzyme-substrate interactions. The final product of this extraction is presented in 6B-D. The same 10 papers were run through Step 4 of the FuncFetch timeline. Code modifications were made to provide timestamps for the beginning and end of processing for each paper. The resulting time intervals were then compared to the human readers. (7B) Reader-1 extracted information. (7C) Reader-2 extracted information. (7D) Reader-3 extracted information. (7B) FuncFetch Step 4 extracted information.

**Supplementary File 8:** Analysis of hallucinations. (8A) First example of messages presenting information from model weights. The pdf method describes searching the document for mentions of the target enzyme families, but only returns entries with the target names and no additional information. In this case, the text method returns a set of correct enzyme activities, but with no relation to information in the text. The information presented is similar to the output when the same model is asked about the enzyme family in question with no added context (see 8C). (8B) Second example of messages presenting information from model weights. In this case, both pdf and text methods present similar information to the output when the same model is asked about the enzyme family in question with no added context (see 8C). (8C) Screenshot of ChatGPT response with GPT-4o model when prompted to describe UDP-glycosyltransferases and their substrates.

